# Deep learning trained on H&E tumor ROIs predicts HER2 status and Trastuzumab treatment response in HER2+ breast cancer

**DOI:** 10.1101/2021.06.14.448356

**Authors:** Saman Farahmand, Aileen I Fernandez, Fahad Shabbir Ahmed, David L. Rimm, Jeffrey H. Chuang, Emily Reisenbichler, Kourosh Zarringhalam

## Abstract

The current standard of care for many patients with HER2-positive breast cancer is neoadjuvant chemotherapy in combination with anti-HER2 agents, based on HER2 amplification as detected by *in situ* hybridization (ISH) or protein immunohistochemistry (IHC). However, hematoxylin & eosin (H&E) tumor stains are more commonly available, and accurate prediction of HER2 status and anti-HER2 treatment response from H&E would reduce costs and increase the speed of treatment selection. Computational algorithms for H&E have been effective in predicting a variety of cancer features and clinical outcomes, including moderate success in predicting HER2 status. In this work, we present a novel convolutional neural network (CNN) approach able to predict HER2 status with increased accuracy over prior methods. We trained a CNN classifier on 188 H&E whole slide images (WSIs) manually annotated for tumor regions of interest (ROIs) by our pathology team. Our classifier achieved an area under the curve (AUC) of 0.90 in cross-validation of slide-level HER2 status and 0.81 on an independent TCGA test set. Within slides, we observed strong agreement between pathologist annotated ROIs and blinded computational predictions of tumor regions / HER2 status. Moreover, we trained our classifier on pre-treatment samples from 187 HER2+ patients that subsequently received trastuzumab therapy. Our classifier achieved an AUC of 0.80 in a five-fold cross validation. Our work provides an H&E-based algorithm that can predict HER2 status and trastuzumab response in breast cancer at an accuracy that is better than IHC and may benefit clinical evaluations.

## Introduction

Human epidermal growth factor 2 (HER2) is a proto-oncogene that is amplified in 20-30% of breast cancer cases (1,2). In the absence of systemic adjuvant therapy, HER2 gene amplification or protein overexpression is associated with aggressive clinical behavior and poor survival outcome (2,3). Fortunately, anti-HER2 treatments such as trastuzumab significantly improve survival outcome (4). Response and overall survival rates of trastuzumab treatment, in combination with chemotherapy, for HER2+ cases for metastatic breast cancer range from 10%-41% and 56-85% respectively, while the response and survival rates for non-metastatic cases range from 50-70% and 56-88% (5–14). As a result, HER2 testing is routinely applied in invasive breast cancer cases as the sole biomarker for anti-HER2 treatment (15,16), however not all clinically defined HER2+ cases respond to treatment nor do tumors lacking HER2 amplification (16). Current ASCO/CAP (16) standards for determining HER2 gene amplification and protein overexpression are *in situ* hybridization (ISH) and immunohistochemistry (IHC) respectively (16–18), though discordance between ISH and IHC is not uncommon and can lead to HER2+ overdiagnosis. One solution may be hematoxylin & eosin (H&E) images, which are commonly generated during pathological analysis and widely abundant, providing opportunities for novel data-driven computational methods. Machine learning-based predictors trained on annotated H&E data could be a potent technology to improve the speed, accuracy, and cost of predicting HER2 status and anti-HER2 treatment response.

In recent years, there has been a growth in machine learning approaches, especially deep learning, in the field of pathology (19). These typically utilize Convolutional Neural Network (CNN) architectures, such as AlexNet (20), GoogleNet (21), or ResNet (22), etc., pre-trained on generic images, and then fine-tune them by re-training the last layers for a specific classification task. This approach is typically referred to as “transfer-learning”. In contrast, the CNN models can be trained using a “full-training” strategy, where no pre-training is utilized, and all CNN parameters are trained using the training dataset of interest.

Representative examples of CNN-based models for pathology applications include tumor/benign classification (23–27), predicting mutations in key genes (24, 27, 28), cancer subtype classification and morphology analysis (23,30), and treatment outcome prediction (31,32). These models have shown impressive performance, demonstrating that subtle molecular features of cancer may be discernible from H&E images.

The objective of this work is to provide a deep learning framework to predict HER2 status and response to trastuzumab therapy from breast cancer H&E slides. Three recent studies have addressed aspects of this problem. Naik, *et al.* (33), developed a ReceptorNet ER+/ER-binary classifier trained on patches sampled from Whole Slide Images (WSIs). They utilized 2535 H&E images from Australian Breast Cancer Tissue Bank (ABCTB) and 1014 H&E images from 939 patients from The Cancer Genomic Atlas (TCGA) with ER, PR, and HER2 status determined by pathologists using IHC. Their classifier achieved an Area Under the Curve (AUC) of 0.899 (95% CI: 0.884–0.913) on cross-validation and an AUC of 0.92 (95% CI: 0.892–0.946) on the test set. They reported that the ER+/ER− classifier performed significantly better on HER2-samples (AUC = 0.927, 95% CI: 0.912–0.943) as compared to HER2 samples (AUC = 0.768, 95% CI: 0.719–0.813). Additionally, they trained and evaluated their classifier to predict PR and HER2 status and obtained an AUC of 0.810 (95% CI: 0.769–0.846) on PR and an AUC of 0.778 (95% CI: 0.730–0.825) on HER2. Rawat *et al.* (34), trained a CNN classifier on 939 TCGA H&E images to predict HER2 status and were able to achieve slide-level AUCs of 0.71 through internal cross-validation, though unexpectedly they observed a higher AUC (0.79) on an independent set (35). Bychkov *et al.* (36), investigated whether predicting HER2 status using a CNN model can guide the choice of therapy. The study utilized cancer tissue samples from FinProg patient series (35), the FinProg validation series (37), and the FinHer clinical trial (38), all of which had HER2 amplification determined by CISH. Their CNN model, trained on 693 H&E-stained patient samples from the FinProg series, was able to predict tile-level HER2 status with AUC 0.70 in a 5-fold cross validation and AUC 0.67 on 712 test images from the FinHer dataset. They also devised a score for HER2 status *(H&E-ERBB2* score) and reported that CISH HER2+ patients with high H&E-*ERBB2* score treated with trastuzumab had a more favorable distant disease-free survival rate than those with a low H&E-*ERBB2* score (Hazard Ratio, 0.37; 95% CI, 0.15–0.93; P = 0.034). CISH HER2+ / high H&E-*ERBB2*-positive patients not treated with trastuzumab also exhibited less favorable disease-free survival (Hazard ratio, 2.03; 95% CI, 0.69–5.94; P = 0.20). These findings indicate that an H&E-based score can contribute to a more accurate prediction of trastuzumab efficacy than CISH alone, but at the same time it is critical to further improve on these AUC values to optimize applicability to clinical practice.

In this work we present novel CNN classifiers for HER2 status and trastuzumab response prediction. We trained a HER2 status predictor model on 188 HER2+/− H&E slides generated from the Yale Pathology electronic database (Yale HER2 cohort). For an independent test set, we utilized 187 HER2+/− H&E slides from the TCGA-BRCA cohort. For trastuzumab response prediction, we used a cohort of 85 pre-treatment HER2+ samples from the Yale Pathology electronic database (Yale Response cohort). In our approach, we employed both transfer and full training strategies. Importantly, we utilized tiles from H&E WSIs, manually annotated for ROIs by our pathology team. We demonstrate that the use of tile-level annotations significantly improves classification accuracy for both HER2 status and trastuzumab response. **Fig. 1** shows an overview of our approach.

**Figure 1.**
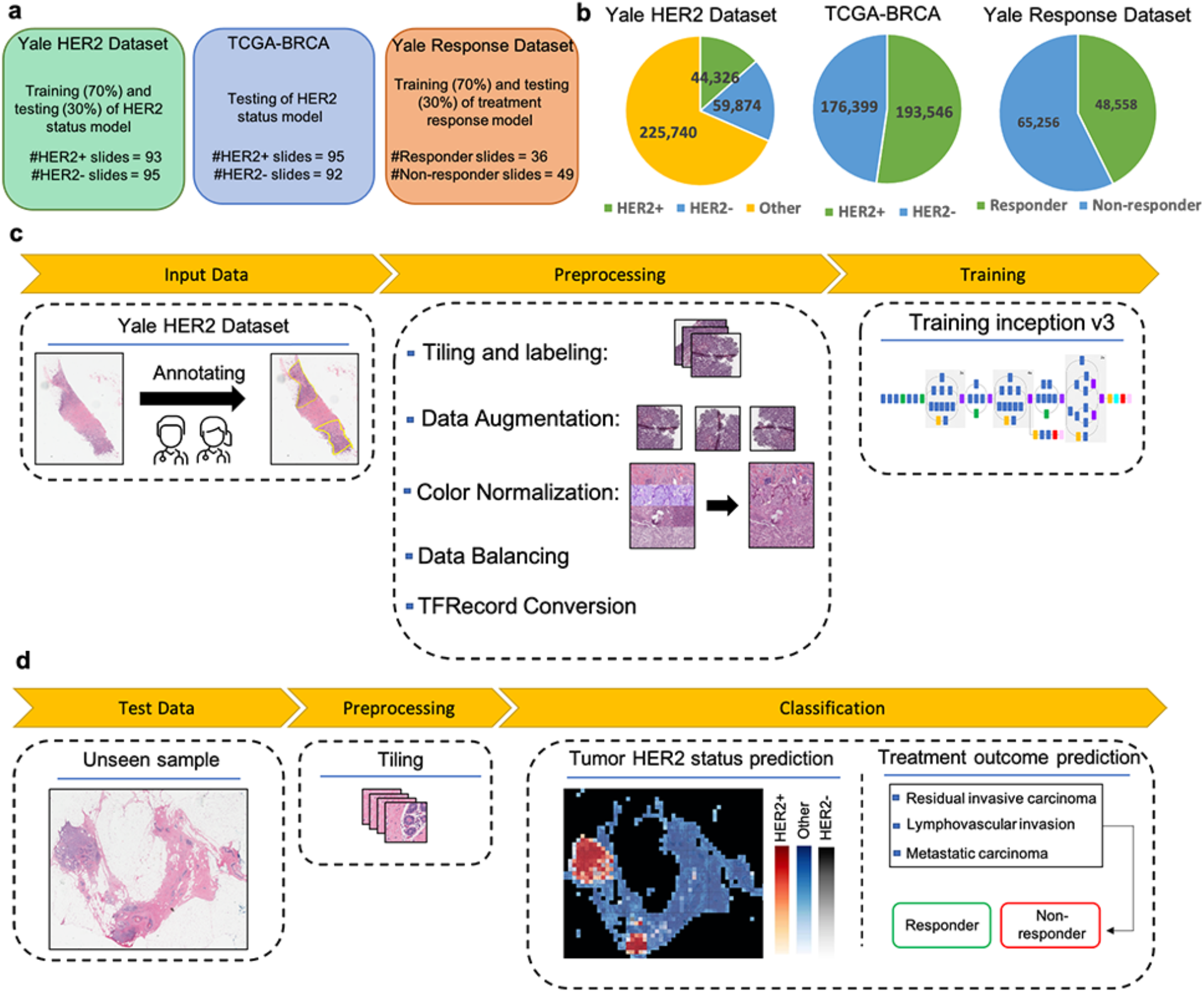
Datasets and study design for HER2 status and Trastuzumab response classification. (a) Datasets generated and used for training and testing the models. (b) Number of tiles in each class. The whole TCGA-BRCA slides as an independent test set were used for testing (we only showed proportion of tiles corresponding to only tumor regions here). For the response model, we only used the tumor regions to train and test the model which the proportion of tiles in each class are depicted here. (c) The main steps for preprocessing of slides and training the model. Our pathology team performed quality checks and annotated the ROIs in every slide to distinguish HER2+ tumor regions, HER2-tumor regions, and other non-tumor regions. In the preprocessing step, slides were tiled into 512×512 pixel windows, and background tiles were removed. Data were randomly split into 70% for training and 30% for testing for both Yale cohorts. The TCGA-BRCA cohort was used to independently validate the HER2 status prediction model. Data augmentation and color normalization were utilized to increase reproducibility. Classes were balanced with down- and up-sampling. Tiles were randomly sorted and converted into TFRecords to train the inception v3-based model. (d) Test data was used to assess model performance. Predictions were visualized on WSIs with heatmaps.

## Materials and Methods

### Data and study design

#### Approval of human tissue data

All tissues and data were retrieved under permission from the Yale Human Investigation Committee protocol #9505008219 to DLR

#### Yale HER2 cohort

188 HER2 positive and negative invasive breast carcinomas were identified by retrospective search of the Yale Pathology electronic database with HER2 positive cases defined as those with 3+ score by immunohistochemistry (IHC) or an equivocal (2+) IHC score with subsequent amplification by fluorescence *in situ* hybridization (FISH) as defined by American Society of Clinical Oncology/College of American Pathologists (ASCO/CAP) clinical practice guidelines (18). H&E slides generated at Yale School of Medicine include 93 HER2+ and 95 HER2-slides. The slides were scanned at Yale Pathology Tissue Services and underwent a slide quality check before they went into the scanner. Broken slides, slides with broken coverslips, and slides with no/minimal tissue were removed. The tissue samples were scanned using Aperio ScanScope Console (v10.2.0.2352) using bright field whole slides scanning at 20× magnification.

#### TCGA HER2 cohort

A total of 668 TCGA-BRCA HER2+/− samples with available HER2 status were downloaded from the GDC portal. Slides were visually inspected by our pathology team to exclude low quality samples with tissue folding or those that appeared to be from frozen tissue. A total of 187 samples (92 HER2- and 95 HER2+) were retained for use as independent test set.

#### Yale trastuzumab response cohort

The response cohort cases were identified also by retrospective search of the Yale Pathology electronic database. Cases included those patients with a pre-treatment breast core biopsy with HER2 positive invasive breast carcinoma who then received neoadjuvant targeted therapy with trastuzumab +/− pertuzumab prior to definitive surgery. HER2 positivity was defined as previously described for the HER2 negative/positive cohort. The response to targeted therapy was obtained from the pathology reports of the surgical resection specimens and dichotomized into responders or non-responders. Those with a complete pathologic response, defined as no residual invasive, lymphovascular invasion or metastatic carcinoma, were designated as responders (n=36). Cases with only residual *in situ* carcinoma were included in the responder category. Those cases with any amount of residual invasive carcinoma, lymphovascular invasion or metastatic carcinoma were categorized as non-responders (n=49).

### Data Preparation

#### Data annotation

Annotation of digital slides was performed, circling areas of invasive carcinoma (Region of Interests, ROIs). Regions of necrosis, *in situ* carcinoma or benign stroma and epithelium were excluded. The images were annotated with ROIs associated to HER2+/− tumor area (TA) by a senior breast pathologist. The annotations were marked tumor boundaries and annotated by Aperio ImageScope software (39). The annotations were exported from the Aperio software in The Extensible Markup Language (XML) format, including X and Y coordinates corresponding to the annotated regions. We used these coordinates for each slide image to tile these regions separately from the rest of the image, labeled as HER2+ or HER2-class.

#### Data prep-processing

Yale cohort slides were randomly split and assigned to 70% for training and 30% for testing. Image slides were tiled into non-overlapping patches of 512 × 512 pixels in 20× magnification. Regions with excess background or containing no tissue as well as regions excess fat were removed as previously described (24). Tiles were shuffled and assigned to Tensorflow Records (TFRecords). To mitigate the effects of class imbalance, we utilized undersampling of the majority class for two-class classification and undersampling and oversampling of the minority and majority classes respectively for three-class classification.

#### Data augmentation and normalization

Data augmentation was performed on training tiles with 90-, 180-, and 270-degree rotations and as well as horizontal and vertical flips. To standardize the color space and address potential batch-effects, we utilized a deep learning-based generative model to normalize the stain color across training and independent test data sets (40). The normalization method is fully unsupervised and does not utilize class label information in the normalization process.

### Model training and assessment

#### CNN architecture and training

We utilized an Inception v3 architecture (40) to predict HER2 status in breast cancer and trastuzumab treatment in HER2+ samples. Models were trained using both transfer learning and full training strategies. Transfer learning model parameters were set according to optimal values from the ImageNet competition (41). The parameters of the last layer of the network were fine-tuned on samples using back propagation. To quantify the impact of ROIs on training, we utilized three different training schemes to predict the HER2 status. **1.** *Unannotated two-way classifier:* Tiles were assigned the label according to the WSI (Positive or Negative) and ROIs were not taken into consideration in training. **2.** *Annotated two-way classifier:* Only tiles falling within the ROIs were utilized in training. Exterior regions (including stromal cells, necrotic cells and/or mixed of tumor and normal cells) were not taken into consideration for training and within ROI tiles were assigned the WSI label. **3.** *Annotated three-way classifier.* Both within ROIs and exterior regions were utilized to train a multi-way classifier. Within ROI tiles were labeled as Positive or Negative according to the WSI label. Tiles in the exterior regions were labeled as “Other” independent of the WSI label. A similar strategy was taken to train a binary classifier for the trastuzumab response predictor. A softmax link function was utilized as loss and predicted probabilities were calculated for each tile. We used RMSProp69 optimization with learning rate of 0.1, weight decay of 0.9, momentum of 0.9, and epsilon of 1.0 to train the model (**Fig. 1**).

#### Model assessment

Model performance was evaluated on test tiles. Slide-level probabilities were calculated by averaging the output probabilities for HER2+ and HER2-classes and the final slide-level label was decided using a 0.5 cutoff threshold on the aggregate probabilities. Model performance was assessed on a per-tile and a per-slide basis. The ROC curves and the corresponding AUC were calculated, and 95% Confidence Intervals (CIs) were estimated by 1,000 iterations of the bootstrap method (42). For the treatment response predictor, we also utilized a 5-fold cross validation, which was possible due to the smaller number of samples. The mean and standard deviation of AUC values were calculated using prediction on each fold.

#### Computational configuration

All analyses were performed in Python. Inception V3 code was adopted from (24). Images were analyzed and processed using OpenSlide. Classification metrics were calculated using the Scikit-learn package (43). All of the computational tasks were performed on Massachusetts Green High Performance Computing Cluster (MGHPCC) on nodes with the following specification: 8 CPUs with 64 GB RAM, Tesla V100 GPUs with 256 GB RAM. TensorFlow and TF-slim documentations and NVIDIA GPUs support were followed to setup and configure CUDA 8.0 Toolkit and cuDNN v5.1.

## Results

### HER2 status classification using unannotated slides

As a base model and for benchmarking with previous approaches, we trained a CNN model to predict HER2 status using unannotated slides from the Yale HER2 cohort (93 HER2+ and 95 HER2-). In this classification scheme (two-way unannotated, Methods) WSIs were tiled to non-overlapping regions and each tile was assigned the label of its corresponding slide (HER2+ or HER2-). The CNN model was trained using both transfer learning as well as full training. Prediction on the left-out data showed a slide-level AUC 0.81 (95% CI, 0.65 – 0.92) in transfer learning and AUC 0.82 (95% CI, 0.66 – 0.92) in the fully trained model. At the tile-level the model achieved an AUC of 0.65 (95% CI, 0.63 – 0.66) in transfer learning and 0.65 (95% CI, 0.63 – 0.65) in the fully trained model (**Fig. 2**). In the following sections, we will present classification schemes that improve on these widely used two-way classifiers, resulting in significant gains in model accuracy and generalizability.

**Figure 2.**
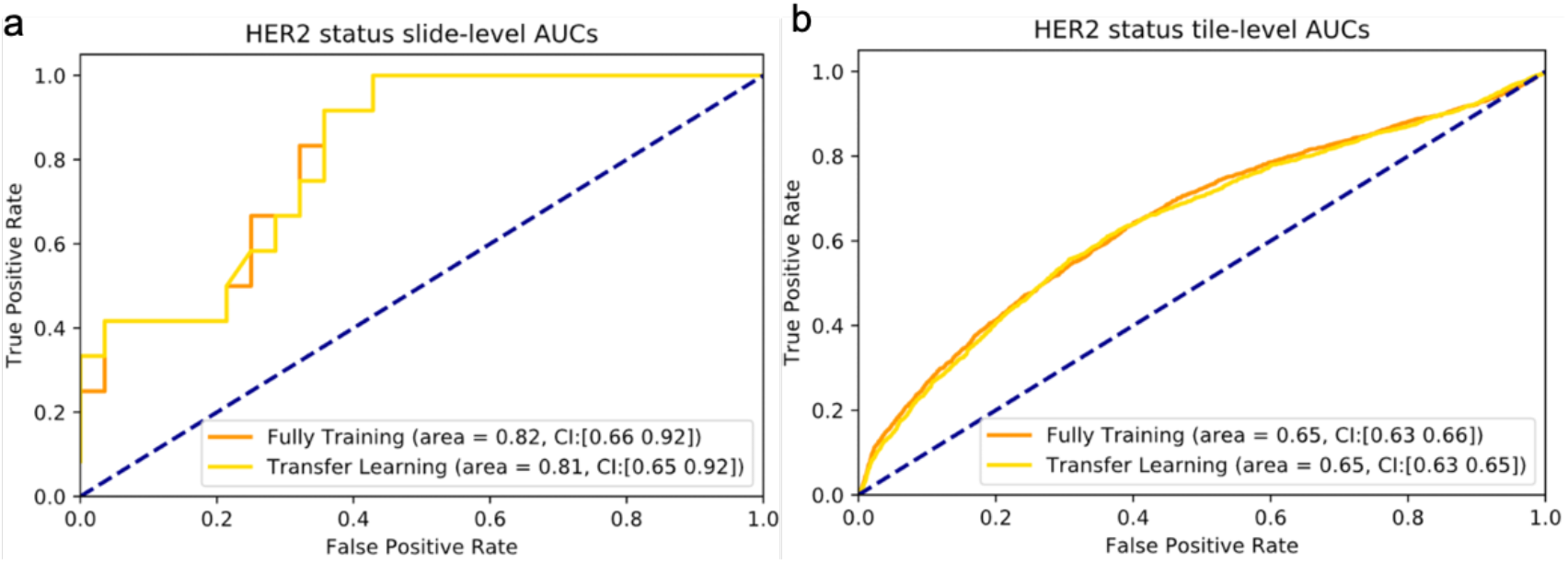
HER2 status classification using unannotated slides. AUC/ROC for HER2 status classification at the slide-level (a) and at the tile-level (b) for both transfer learning and the fully trained models.

### HER2 status classification using annotated slides

We hypothesized that using tiles from ROIs may reduce irrelevant features and enable the CNN to better learn features specific to HER2+/− tumor status. Our pathology team annotated the Yale HER2 cohort to mark the regions corresponding to the invasive tumor cells while excluding regions such as necrosis, *in situ* carcinoma, benign stroma and epithelium. Each slide was masked according to these manual annotations, then broken into tiles for analysis with each tile categorized as tumor or Other. The tumor tiles were categorized as either HER2+ or HER2-, yielding 3 classes for training: HER2+, HER2- and Other. To draw direct comparison with the previous classifier, we trained another two-way classifier, this time trained on HER2+/− tiles only (two-way annotated, Methods). **Fig. 3.a** and **b** present the AUC values of the CNN classifier for the two-way annotated model using both full training and transfer learning approaches. The model achieved a slide-level AUC of 0.90 (95% CI, 0.79 – 0.97) and a tile-level AUC of 0.77 (95% CI, 0.76 – 0.77) in the transfer learning approach, and AUC of 0.89 (95% CI, 0.78 – 0.96) and a tile-level AUC of 0.75 (95% CI, 0.74 – 0.75) in the fully trained model. We generated a heatmap using tile-predicted probabilities to visualize the predictions made by the model (**Fig. 3.d** middle column). Although the prediction performance at the slide-level is high, the tile-level heatmaps do not show the same level of performance compared with tile-level pathologist annotations (**Fig. 3.d** first column yellow regions).

**Figure 3.**
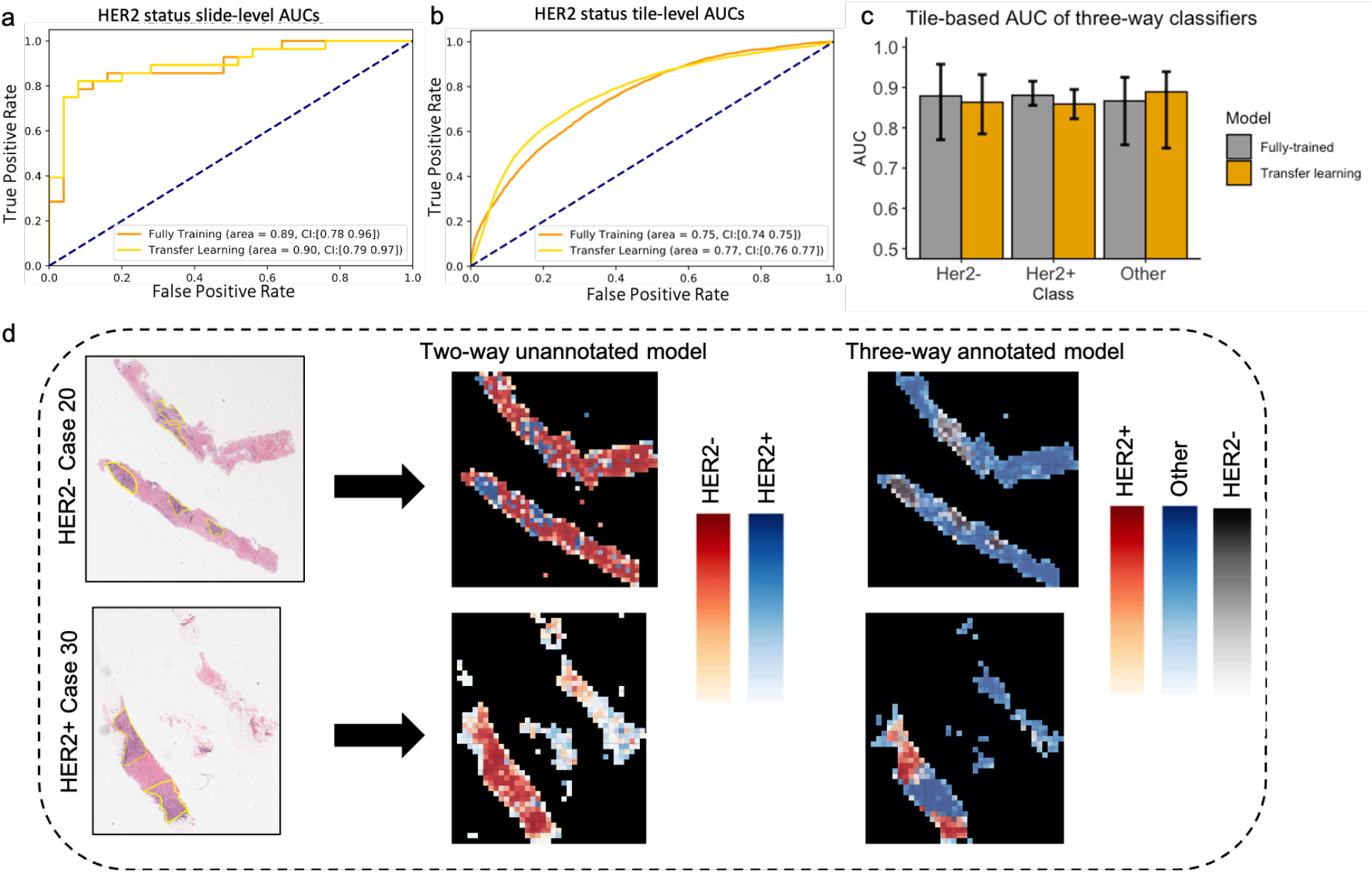
HER2 tumor status classification using annotated slides. AUC/ROC for HER2 status classification at the slide-level (a) and at the tile-level (b) for both transfer learning and the fully trained models. (c) Tile-level AUC of the three-way classifiers. (d) Two representative H&E slides from the test set with corresponding heatmaps based on predicted probabilities by the CNN model. Yellow regions indicate the ROIs determined by our pathology team. The middle panel shows predicted heatmaps for unannotated model and the right panel shows the predicted heatmaps from three-way annotated model.

Next, we tested whether including tiles from exterior regions of ROIs can improve the tile-level accuracies and ROI visualizations by heatmaps. Our rationale for including these tiles was that the classifier trained on HER2+/− tiles is likely unable to predict class label of the exterior tiles. We trained a three-way classifier on HER2+, HER2-, and Other tiles (annotated three-way, Methods). **Fig. 3.c** shows the tile-level AUCs for models trained using transfer learning or full training. The fully-trained CNN model predicted the HER2-status with a AUC of 0.88 (95% CI, 0.77 – 0.95), and HER2+ with an AUC of 0.88 (95% CI, 0.85 – 0.91) and Other class with an AUC of 0.87 (95% CI, 0.75 – 0.92). The transfer learning model achieved similar AUCs. This is a clear increase in the CNN’s tile-level AUC compared with the two-way annotated classifier (**Fig 3b**), indicating that features from non-HER2+/− tiles can decrease the confusion between HER2+ and HER2-tiles. **Fig. 3.d** right column illustrates the heatmaps produced by the three-way classifier. There is strong agreement between the heatmap from the three-way classifier and pathologist annotated ROIs, indicating the utility of our model for automatic ROI detection.

### Model validation on independent test set

We next validated the HER2 status classifier on an external independent test set. For this analysis, we downloaded a dataset consisting of 569 HER2- and 99 HER2+ WSIs of H&E stained sections of formalin fixed paraffin embedded (FFPE) samples from TCGA-BRCA cohort. Our pathology team performed quality control to exclude samples with poor scanning and staining quality, resulting in 197 samples being excluded from further analysis. Slides were processed and tiled as in the Yale HER2 cohort, resulting in 176399 HER2+. and 193546 HER2-tiles. Since the training and test cohorts were from independent sources, we performed a stain-color normalization step using a deep generative model (40) to scale the TCGA cohort to the Yale cohort (**Methods**). The CNN HER2 status classifier, trained on the Yale HER2 cohort was used to make predictions on the TCGA cohort. The AUCs of the model performance are 0.81 (95% CI: 0.73 – 0.84) at the slide-level and 0.65 (95% CI: 0.54 – 0.69) at the tile-level.

We also tested whether ROIs can be accurately detected in the test set. As in the Yale HER2 cohort, our pathology team annotated the TCGA-BRCA cohorts to mark ROIs. **Fig. 4** shows two representative samples of annotations (yellow regions), heatmaps predicted by the unannotated two-way classifier, and the heatmap produced by the annotated three-way classifier. There is a high-level of agreement between the ROIs and the predictions made by the three-way classifier, demonstrating the generalizability of our automatic ROI detection.

**Figure 4.**
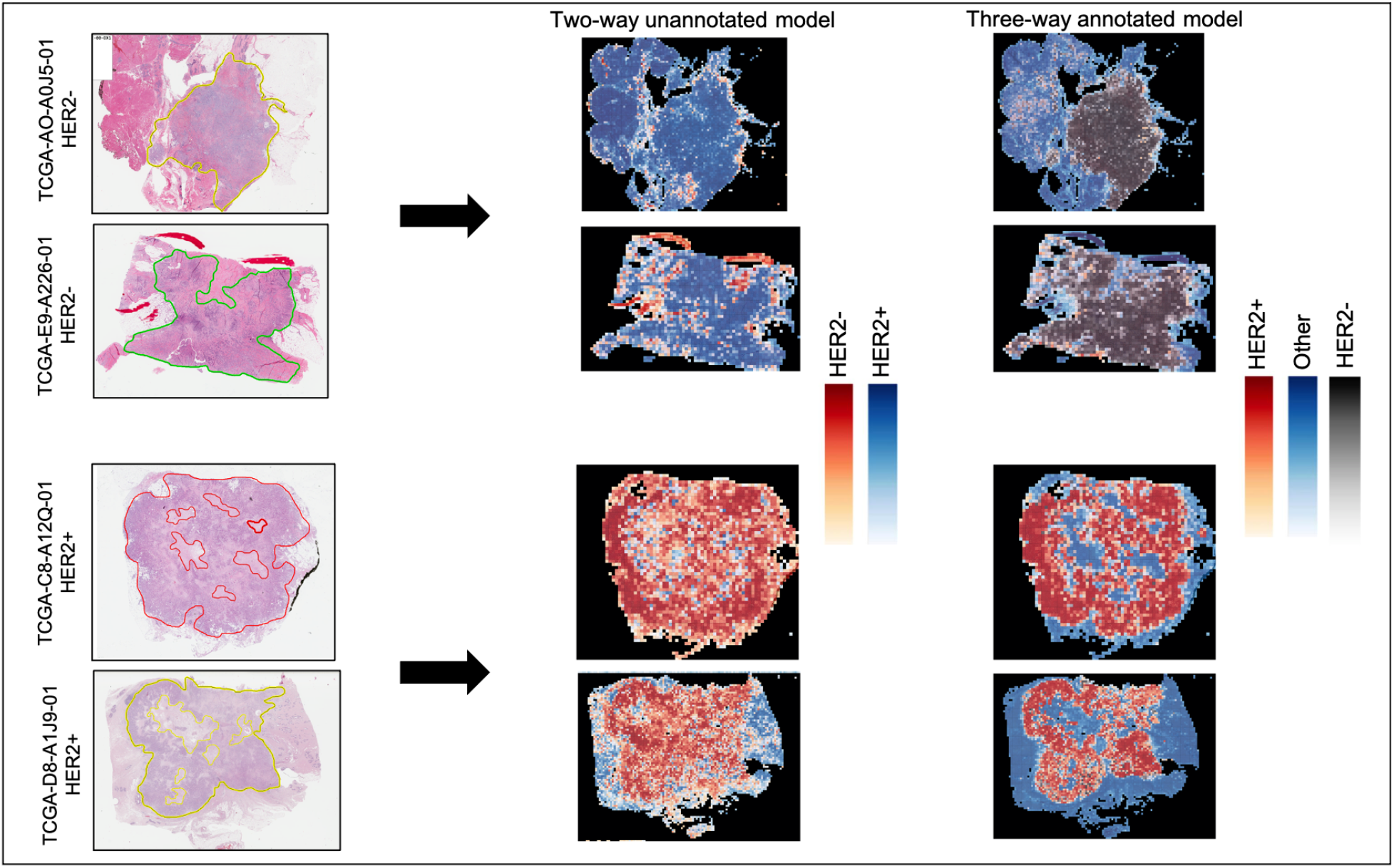
Representative H&E slides from TCGA test set and their predicted heatmaps. Left column slides along with indicated ROIs. Middle panel: Two-way unannotated classifier heatmaps. Right panel: Three-way annotated classifier heatmaps.

### Deep learning predicts trastuzumab treatment outcome

We next tested whether deep learning trained on H&E slides from HER2+ patients can predict trastuzumab treatment outcome. For this study we utilized pre-treatment H&E slides from the Yale trastuzumab cohort. As with the previous samples, our pathology team annotated the slides to mark the invasive tumor cells area. In addition to model assessment using test sets and CI estimation with bootstrapping, we performed a 5-fold cross validation to more stringently assess model performance. The unannotated model achieved an AUC of 0.68 (95% CI: 0.47 – 0.88) at the slide-level and an AUC of 0.63 (95% CI: 0.62 – 0.65) at the tile-level. On the other hand, the annotated models achieved an AUC of 0.80 (95% CI: 0.69 – 0.88) at the slide-level and an AUC of 0.73 (95% CI: 0.63 – 0.79) at the tile level (**Fig 5**). As in the HER2 status classifier, the improvement in AUCs shows the importance of annotations in training of deep learning classifiers for response prediction.

**Figure 5.**
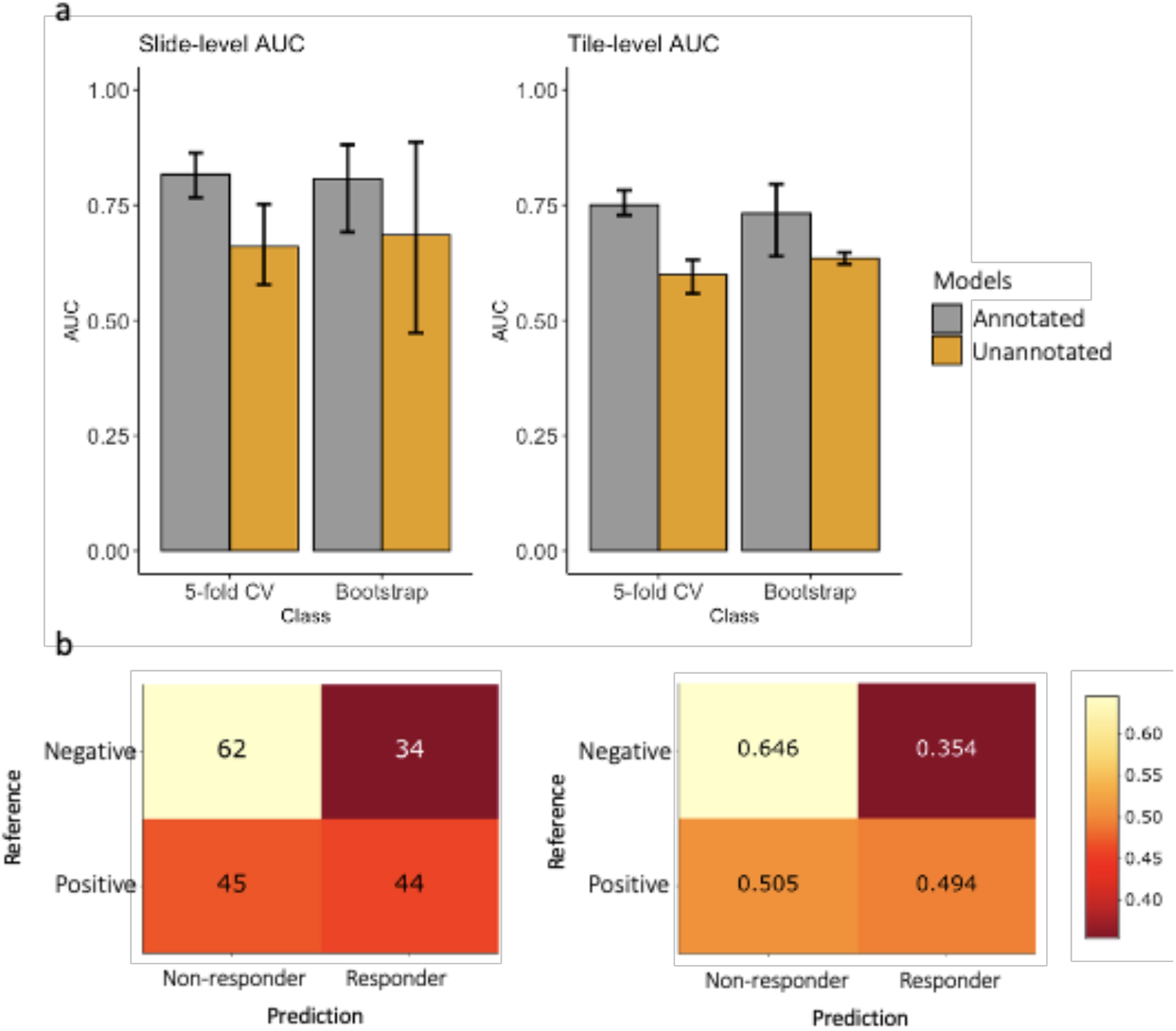
Trastuzumab response prediction. **(a)** Slide-level and Tile-level AUC/ROC for both annotated and unannotated models using bootstrapping and 5-fold cross validation. (b) Confusion matrix of the HER2 status classifier used as Trastuzumab response predictor.

We also tested whether the HER2 status classifier can directly predict trastuzumab treatment response. This is important as HER2 status is the clinical biomarker for anti-HER2 treatment. For this test, the Yale HER2 response data was used as input to the HER2 status CNN-classifier, and the predictions made by the classifier (HER2+/−) were tabulated against the response labels. **Fig. 5.b** shows the confusion matrix. As shown, 50% of HER2+ samples were predicted as responders, while 65% of the HER2-samples were predicted as non-responders. These results demonstrate that although HER2 status, as determined by traditional IHC/ISH methods, can moderately predict trastuzumab response, more specifically trained models are needed to better identify patients who would benefit from trastuzumab treatment therapy. Taken together, these results support the feasibility of image-based biomarkers for prediction of trastuzumab therapy and the ability of the deep learning model to identify morphological variations associated with treatment outcome. Trastuzumab response predictors, such as the one presented in this study, have the potential to augment HER2 status testing for treatment recommendation in HER2+ patients.

## Discussion

In this work we presented CNN-based classifiers for determining HER2 status and trastuzumab response prediction. Using high-quality slides carefully annotated by expert pathologists, we were able to reduce the feature space and hence the number of required samples for training our classifiers. A two-way classifier of HER2 status, trained on within-tumor ROI tiles achieved a slide-level AUC of 0.90 in cross-validation and 0.81 on an independent test set. To increase the tile-level accuracy and the predicted ROI heatmaps, we devised a three-way classification scheme and trained a multi-way classifier using tiles from within as well as the exterior ROI regions. The three-way classifier was able to distinguish tiles from each class with high accuracy (AUCs: HER2+ 0.88, HER2-0.88, Other: 0.87). Heatmaps produced by the three-way classifier show a remarkable agreement with pathology annotations, both in the slides from the training set as well as the slides from the independent set.

In a recent study, Bychkov *et al.* (36) trained a two-way CNN model to predict HER2 status on H&E slides from FinProg patient series. Their classifier was trained on random tile crops of size 950×950 from 3500×3500 WSIs and achieved a tile-level AUC of 0.70 (95% CI, 0.63-0.77) in cross validation and AUC of 0.67 (95% CI, 0.62-0.71) on test images from the FinHer dataset. They did not report slide-level AUCs. In their approaches, only tiles from the center crop (2100 × 2100 pixels) of the WSI were used to test the prediction performance, whereas in our approach tile-level AUC was estimated using all test tiles. As such, direct comparison between the methods is confounded by their test-tile selection procedure. On the other hand, our ensemble procedure for slide-level HER2 status prediction results in a significant increase in AUCs, demonstrating the robustness and the generalizability of our approach.

In another related study Rawat *et al.* (34) trained a patch-based CNN classifier on a TCGA cohort with patch-sizes of 224 × 224. Their model achieved a slide-level HER2 AUC of 0.71 (TCGA, n = 124) in a 5-fold cross validation. They also tested the generalizability of their model using an independent cohort from The Australian Breast Cancer Tissue Bank (ABCTB) (35). Their model achieved a slide-level Her2 AUC = 0.79 (ABCTB, n = 2487). Interestingly, this AUC is larger than their within-TCGA cross-validation AUC (0.71), on which their model was trained, presumably due to the higher quality of ABCTB slides. In both cases, our cross-validation and independent test AUC on TCGA cohort (0.9, and 0.81) improves upon these results. Finally, Naik, et al Naik, *et al.* (33), trained a receptor net classifier on ABCTA cohort that achieved a slide-level AUC of 0.778 (95% CI: 0.730– 0.825) on HER2 status.

A key improvement of our method compared to these previous approaches is our use of tumor Regions of Interest (ROI) annotations during training. These annotations allowed us to train and evaluate the three-way classification model for HER2+, HER2-, and non-tumor tiles within each WSI. In contrast, previous approaches have utilized a weakly supervised two-way (HER2+/HER2-) classification model based on slide-level rather than tile level annotations.

Another strength of our method is the use of deep learning-based color normalization (40) to remove batch-effects and improve generalizability to independent datasets. We utilized a deep learning-based color normalization scheme developed by Zanjani *et. al* (40). Color and intensity variations between H&E samples from different medical centers or even within the same laboratory samples generated at various trials or time periods is common (44). Variations in specimen sample preparation protocol, staining protocols, scanning, and imaging device characteristics are some of the contributing factors. As such, H&E stain color normalization has been studied and used in deep learning approaches (45–47). Recently (44), Howard *et. al* showed that features extracted by deep learning models trained on H&E images vary substantially across data sets. They point out that color normalization alone may not be sufficient to address confounding factors and generalizability of deep learning models to independent datasets remains a challenging task. However, this may be limited to more subtle molecular features of cancer. In our case, color normalization resulted in a small increase in model accuracy, and further investigation of similar effect are likely important to understanding the variations in predictive accuracy across different cohorts.

Taken together, the significant improvement in slide-level and tile-level AUCs relative to those from our unannotated model and previous results (33,34,36) indicate the importance of using pathology annotation to guide targeted feature learning.

Although the response rate to trastuzumab therapy in HER2+ patients has been good, augmenting HER2 status determination with more accurate methodologies for treatment response prediction has the potential to improve patient care. Using pre-treatment samples from HER2+ positive patients with known trastuzumab response, we trained a classifier able to accurately predict response (AUC: 0.80; 5-fold cross validation). In contrast, Bychkov *et. al* (36) showed that their HER2 status score was associated with survival hazard ratio on a trastuzumab treated cohort. That approach, while conceptually informative, lacks the direct clinical applicability of a binary response predictor as we have presented in this study. Indeed, we showed that the HER2 status classifier is a weak predictor of trastuzumab response, demonstrating the value of directly predicting anti-HER2 response efficacy and suggesting the need for further biomarker investigation.

In summary, the methodology that we have developed in this study provides an accurate and reproducible H&E-based approach for detection of HER2 status and response to trastuzumab therapy. Given that many new drugs have emerged for treatment of patients that express HER2, a combination of an AI classifier with conventional methods might improve the ability to select which HER2 drug is most likely to benefit each patient. Future prospective trials in the neoadjuvant setting are being considered. Furthermore, we anticipate that this approach will be generalizable to other cancer types and treatment outcome predictions.

## Data/Code Availability

Data and code are available upon request.

## Acknowledgements

J.H.C. acknowledges support from NCI grant R01CA230031.

E.S.R. acknowledges support from Grant #IRG 17-172-57 from the American Cancer Society.

D.L.R. acknowledges support from the Breast Cancer Research Foundation BCRF20-138

## Author Contributions

S.F. developed the classifiers, analyzed the Yale and TCGA data, and drafted the paper. A.F. performed quality control and analyzed the Yale data. F.S.A performed quality control and analyzed the Yale data. D.R. provided pathological evaluations and oversaw the project. J.H.C oversaw the project and drafted the paper. E.S.R generated the data, annotated the data, provided pathological evaluations and oversaw the project. K.Z led the project and finalized the paper.

## Supplementary information is available at Modern Pathology’s website

## References

1. Slamon DJ, Clark GM, Wong SG, Levin WJ, Ullrich A, McGuire WL. Human breast cancer: correlation of relapse and survival with amplification of the {HER-2/neu} oncogene. Science (80-). 1987 Jan;235(4785):177–82.

2. Press MF, Bernstein L, Thomas PA, Meisner LF, Zhou JY, Ma Y, et al. {HER-2/neu} gene amplification characterized by fluorescence in situ hybridization: poor prognosis in node-negative breast carcinomas. J Clin Oncol. 1997 Aug;15(8):2894–904.

3. Tandon AK, Clark GM, Chamness GC, Ullrich A, McGuire WL. {HER-2/neu} oncogene protein and prognosis in breast cancer. J Clin Oncol. 1989 Aug;7(8):1120–8.

4. Hayes DF. {HER2} and Breast Cancer – A Phenomenal Success Story. N Engl J Med. 2019 Sep;381(13):1284–6.

5. Andersson M, Lidbrink E, Bjerre K, Wist E, Enevoldsen K, Jensen AB, et al. Phase {III} Randomized Study Comparing Docetaxel Plus Trastuzumab With Vinorelbine Plus Trastuzumab As {First-Line}Therapy of Metastatic or Locally Advanced Human Epidermal Growth Factor Receptor 2--Positive Breast Cancer: The {HERNATA} Study. J Clin Orthod. 2011 Jan;29(3):264–71.

6. Pivot X, Manikhas A, Żurawski B, Chmielowska E, Karaszewska B, Allerton R, et al. {CEREBEL} ({EGF111438)}: A Phase {III}, Randomized, {Open-Label} Study of Lapatinib Plus Capecitabine Versus Trastuzumab Plus Capecitabine in Patients With Human Epidermal Growth Factor Receptor 2--Positive Metastatic Breast Cancer. J Clin Orthod. 2015 May;33(14):1564–73.

7. Valero V, Forbes J, Pegram MD, Pienkowski T, Eiermann W, von Minckwitz G, et al. Multicenter phase {III} randomized trial comparing docetaxel and trastuzumab with docetaxel, carboplatin, and trastuzumab as first-line chemotherapy for patients with {HER2-gene-amplified} metastatic breast cancer ({BCIRG} 007 study): two highly active th. J Clin Oncol. 2011 Jan;29(2):149–56.

8. Slamon DJ, Leyland-Jones B, Shak S, Fuchs H, Paton V, Bajamonde A, et al. Use of chemotherapy plus a monoclonal antibody against {HER2} for metastatic breast cancer that overexpresses {HER2}. N Engl J Med. 2001 Mar;344(11):783–92.

9. Rugo HS, Barve A, Waller CF, Hernandez-Bronchud M, Herson J, Yuan J, et al. Effect of a Proposed Trastuzumab Biosimilar Compared With Trastuzumab on Overall Response Rate in Patients With {ERBB2} ({HER2)-Positive} Metastatic Breast Cancer: A Randomized Clinical Trial. JAMA. 2017 Jan;317(1):37–47.

10. Urruticoechea A, Rizwanullah M, Im S-A, Ruiz ACS, Láng I, Tomasello G, et al. Randomized Phase {III} Trial of Trastuzumab Plus Capecitabine With or Without Pertuzumab in Patients With Human Epidermal Growth Factor Receptor 2-Positive Metastatic Breast Cancer Who Experienced Disease Progression During or After {Trastuzumab-Based} Th. J Clin Oncol. 2017 Sep;35(26):3030–8.

11. Gianni L, Romieu GH, Lichinitser M, Serrano S V, Mansutti M, Pivot X, et al. {AVEREL}: a randomized phase {III} Trial evaluating bevacizumab in combination with docetaxel and trastuzumab as first-line therapy for {HER2-positive} locally recurrent/metastatic breast cancer. J Clin Oncol. 2013 May;31(14):1719–25.

12. Baselga J, Cortés J, Kim S-B, Im S-A, Hegg R, Im Y-H, et al. Pertuzumab plus trastuzumab plus docetaxel for metastatic breast cancer. N Engl J Med. 2012 Jan;366(2):109–19.

13. Gelmon KA, Boyle FM, Kaufman B, Huntsman DG, Manikhas A, Di Leo A, et al. Lapatinib or Trastuzumab Plus Taxane Therapy for Human Epidermal Growth Factor Receptor 2--Positive Advanced Breast Cancer: Final Results of {NCIC}{CTG}{MA.31}. J Clin Orthod. 2015 May;33(14):1574–83.

14. Blackwell KL, Burstein HJ, Storniolo AM, Rugo H, Sledge G, Koehler M, et al. Randomized study of Lapatinib alone or in combination with trastuzumab in women with {ErbB2-positive}, trastuzumab-refractory metastatic breast cancer. J Clin Oncol. 2010 Mar;28(7):1124–30.

15. Advani PP, Crozier JA, Perez EA. {HER2} testing and its predictive utility in {anti-HER2} breast cancer therapy. Biomark Med. 2015;9(1):35–49.

16. Woo JW, Lee K, Chung YR, Jang MH, Ahn S, Park SY. The updated 2018 American Society of Clinical {Oncology/College} of American Pathologists guideline on human epidermal growth factor receptor 2 interpretation in breast cancer: comparison with previous guidelines and clinical significance of the proposed. Hum Pathol. 2020 Apr;98:10–21.

17. Wolff AC, Hammond MEH, Schwartz JN, Hagerty KL, Allred DC, Cote RJ, et al. American Society of Clinical {Oncology/College} of American Pathologists guideline recommendations for human epidermal growth factor receptor 2 testing in breast cancer. Arch Pathol Lab Med. 2007;131(1):18–43.

18. Wolff AC, Hammond MEH, Allison KH, Harvey BE, Mangu PB, Bartlett JMS, et al. Human Epidermal Growth Factor Receptor 2 Testing in Breast Cancer: American Society of Clinical {Oncology/College} of American Pathologists Clinical Practice Guideline Focused Update. Arch Pathol Lab Med. 2018 Nov;142(11):1364–82.

19. Jiang Y, Yang M, Wang S, Li X, Sun Y. Emerging role of deep learning-based artificial intelligence in tumor pathology. Cancer Commun. 2020 Apr;40(4):154–66.

20. Krizhevsky A, Sutskever I, Hinton GE. {ImageNet} Classification with Deep Convolutional Neural Networks. Commun ACM. 2017 May;60(6):84–90.

21. Szegedy C, Wei Liu, Yangqing Jia, Sermanet P, Reed S, Anguelov D, et al. Going deeper with convolutions. In: 2015 IEEE Conference on Computer Vision and Pattern Recognition (CVPR). 2015. p. 1–9.

22. He K, Zhang X, Ren S, Sun J. Deep Residual Learning for Image Recognition. In: 2016 IEEE Conference on Computer Vision and Pattern Recognition (CVPR). 2016. p. 770–8.

23. Noorbakhsh J, Farahmand S, Foroughi pour A, Namburi S, Caruana D, Rimm D, et al. Deep learning-based cross-classifications reveal conserved spatial behaviors within tumor histological images. Nat Commun [Internet]. 2020 Dec 1 [cited 2021 Mar 14];11(1):1–14. Available from: https://doi.org/10.1038/s41467-020-20030-5

24. Coudray N, Ocampo PS, Sakellaropoulos T, Narula N, Snuderl M, Fenyö D, et al. Classification and mutation prediction from non–small cell lung cancer histopathology images using deep learning. Nat Med [Internet]. 2018 Oct 1 [cited 2020 Nov 13];24(10):1559–67. Available from: https://pubmed.ncbi.nlm.nih.gov/30224757/

25. Wei JW, Tafe LJ, Linnik YA, Vaickus LJ, Tomita N, Hassanpour S. Pathologist-level classification of histologic patterns on resected lung adenocarcinoma slides with deep neural networks. Sci Rep [Internet]. 2019 Dec 1 [cited 2021 Jun 12];9(1):1–8. Available from: https://doi.org/10.1038/s41598-019-40041-7

26. Liu Y, Gadepalli K, Norouzi M, Dahl GE, Kohlberger T, Boyko A, et al. Detecting Cancer Metastases on Gigapixel Pathology Images. 2017 Mar 3 [cited 2021 Jun 12]; Available from: http://arxiv.org/abs/1703.02442

27. Noorbakhsh J, Farahmand S, Soltanieh-Ha M, Namburi S, Zarringhalam K, Chuang J. Pan-cancer classifications of tumor histological images using deep learning [Internet]. bioRxiv. bioRxiv; 2019 [cited 2021 Mar 14]. p. 715656. Available from: https://doi.org/10.1101/715656

28. Coudray N, Tsirigos A. Deep learning links histology, molecular signatures and prognosis in cancer. Nat Cancer [Internet]. 2020 Aug 27 [cited 2021 Jun 12];1(8):755–7. Available from: https://www.nature.com/articles/s43018-020-0099-2

29. Yang Y, Fang Q, Shen H-B. Predicting gene regulatory interactions based on spatial gene expression data and deep learning. Krishnaswamy S, editor. PLOS Comput Biol [Internet]. 2019 Sep 17 [cited 2021 Feb 22];15(9):e1007324. Available from: https://dx.plos.org/10.1371/journal.pcbi.1007324

30. Yu K-H, Wang F, Berry G, Ré C, Altman R, Snyder M, et al. Classifying Non-Small Cell Lung Cancer Histopathology Types and Transcriptomic Subtypes using Convolutional Neural Networks. bioRxiv [Internet]. 2019 Jan 25 [cited 2020 Nov 13];530360. Available from: https://doi.org/10.1101/530360

31. Mobadersany P, Yousefi S, Amgad M, Gutman DA, Barnholtz-Sloan JS, Velázquez Vega JE, et al. Predicting cancer outcomes from histology and genomics using convolutional networks. Proc Natl Acad Sci [Internet]. 2018;115(13):E2970--E2979. Available from: https://www.pnas.org/content/115/13/E2970

32. Braman N, Adoui M El, Vulchi M, Turk P, Etesami M, Fu P, et al. Deep learning-based prediction of response to HER2-targeted neoadjuvant chemotherapy from pre-treatment dynamic breast MRI: A multi-institutional validation study. 2020 Jan 22 [cited 2021 Jun 12]; Available from: http://arxiv.org/abs/2001.08570

33. Naik N, Madani A, Esteva A, Keskar NS, Press MF, Ruderman D, et al. Deep learning-enabled breast cancer hormonal receptor status determination from base-level H&E stains. Nat Commun [Internet]. 2020 Dec 1 [cited 2021 Jun 12];11(1):1–8. Available from: https://doi.org/10.1038/s41467-020-19334-3

34. Rawat RR, Ortega I, Roy P, Sha F, Shibata D, Ruderman D, et al. Deep learned tissue “fingerprints” classify breast cancers by ER/PR/Her2 status from H&E images. Sci Rep [Internet]. 2020 Dec 1 [cited 2021 Jun 12];10(1):1–13. Available from: https://doi.org/10.1038/s41598-020-64156-4

35. Carpenter JE, Marsh D, Mariasegaram M, Clarke CL. The Australian Breast Cancer Tissue Bank (ABCTB). Open J Bioresour [Internet]. 2014 Jul 24 [cited 2021 Jun 12];1(0):e1. Available from: http://dx.doi.org/10.5334/ojb.aa

36. Bychkov D, Linder N, Tiulpin A, Kücükel H, Lundin M, Nordling S, et al. Deep learning identifies morphological features in breast cancer predictive of cancer {ERBB2} status and trastuzumab treatment efficacy. Sci Rep. 2021 Feb;11(1):4037.

37. Lundin J, Lundin M, Isola J, Joensuu H. A web-based system for individualised survival estimation in breast cancer. BMJ [Internet]. 2003 [cited 2021 Jun 12];326(7379):29. Available from: https://pubmed.ncbi.nlm.nih.gov/12511459/

38. Joensuu H, Kellokumpu-Lehtinen P-L, Bono P, Alanko T, Kataja V, Asola R, et al. Adjuvant Docetaxel or Vinorelbine with or without Trastuzumab for Breast Cancer. N Engl J Med [Internet]. 2006 Feb 23 [cited 2021 Jun 12];354(8):809–20. Available from: www.nejm.org

39. Aperio ImageScope – Pathology Slide Viewing Software: Leica Biosystems [Internet]. [cited 2021 Mar 14]. Available from: https://www.leicabiosystems.com/digital-pathology/manage/aperio-imagescope/

40. Zanjani FG, Zinger S, Bejnordi BE, van der Laak JA, de With PHN. Histopathology {Stain-Color} Normalization Using Deep Generative Models. 2018 Apr.

41. Szegedy C, Liu W, Jia Y, Sermanet P, Reed S, Anguelov D, et al. Going deeper with convolutions. In: Proceedings of the IEEE Computer Society Conference on Computer Vision and Pattern Recognition. IEEE Computer Society; 2015. p. 1–9.

42. Efron B, Tibshirani RJ, Cox DR, Reid N, Isham V, Louis TA, et al. An Introduction to the Bootstrap. 1st ed. Taylor & Francis Group; 1994.

43. Pedregosa F, Varoquaux G, Gramfort A, others. Scikit-learn: Machine learning in Python. Mach Learn …. 2011;

44. Howard FM, Dolezal J, Kochanny S, Schulte J, Chen H, Heij L, et al. The Impact of Digital Histopathology Batch Effect on Deep Learning Model Accuracy and Bias. bioRxiv [Internet]. 2020 Dec 4 [cited 2021 Jun 12];2020.12.03.410845. Available from: https://doi.org/10.1101/2020.12.03.410845

45. Wang YY, Chang SC, Wu LW, Tsai ST, Sun YN. A color-based approach for automated segmentation in tumor tissue classification. In: Annual International Conference of the IEEE Engineering in Medicine and Biology – Proceedings [Internet]. Annu Int Conf IEEE Eng Med Biol Soc; 2007 [cited 2021 Feb 7]. p. 6576–9. Available from: https://pubmed.ncbi.nlm.nih.gov/18003532/

46. Alsubaie N, Trahearn N, Raza SEA, Snead D, Rajpoot NM. Stain Deconvolution Using Statistical Analysis of {Multi-Resolution} Stain Colour Representation. PLoS One. 2017 Jan;12(1):e0169875.

47. Sha L, Schonfeld D, Sethi A. Color normalization of histology slides using graph regularized sparse NMF. In: Gurcan MN, Tomaszewski JE, editors. Medical Imaging 2017: Digital Pathology [Internet]. SPIE; 2017 [cited 2021 Feb 7]. p. 1014010. Available from: http://proceedings.spiedigitallibrary.org/proceeding.aspx?doi=10.1117/12.2254214

